# A cascade of destabilizations: combining *Wolbachia* and Allee effects to eradicate insect pests

**DOI:** 10.1101/064774

**Authors:** Julie C. Blackwood, Roger Vargas, Xavier Fauvergue

**Affiliations:** Department of Mathematics and Statistics, Williams College, Williamstown, MA 01267, USA; INRA, CNRS, Université Nice Côte d’Azur, ISA, France

**Keywords:** biological control, cytoplasmic incompatibility, eradication, extinction, mating disruption, transient dynamics

## Abstract

1. The management of insect pests has long been dominated by the use of chemical insecticides, with the aim of instantaneously killing enough individuals to limit their damage. To minimize unwanted consequences, environmentally-friendly approaches have been proposed that utilize biological control and take advantage of intrinsic demographic processes to reduce pest populations.
2. We address the feasibility of a novel pest management strategy based on the release of insects infected with *Wolbachia*, which causes cytoplasmic incompatibilities in its host population, into a population with a pre-existing Allee effect. We hypothesize that the transient decline in population size caused by a successful invasion of *Wolbachia* can bring the population below its Allee threshold and, consequently, trigger extinction.
3. We develop a stochastic population model that accounts for *Wolbachia*-induced cytoplasmic incompatibilities in addition to an Allee effect arising from mating failures at low population densities. Using our model, we identify conditions under which cytoplasmic incompatibilities and Allee effects successfully interact to drive insect pest populations toward extinction. Based on our results, we delineate control strategies based on introductions of *Wolbachia*-infected insects.
4. We extend this analysis to evaluate control strategies that implement successive introductions of two incompatible *Wolbachia* strains. Additionally, we consider methods that combine *Wolbachia* invasion with mating disruption tactics to enhance the pre-existing Allee effect.
5. We demonstrate that Wolbachia-induced cytoplasmic incompatibility and the Allee effect act independently from one another: the Allee effect does not modify the *Wolbachia*-invasion threshold, and cytoplasmic incompatibilities only have a marginal effect on the Allee threshold. However, the interaction of these two processes can drive even large populations to extinction. The success of this method can be amplified by the introduction of multiple *Wolbachia* cytotypes as well as the addition of mating disruption.
6. Our study extends the existing literature by proposing the use of *Wolbachia* introductions to capitalize on pre-existing Allee effects and consequently eradicate insect pests. More generally, it highlights the importance of transient dynamics, and the relevance of manipulating a cascade of destabilizatons for pest management.

## Introduction

Although most insect species provide crucial ecosystem services (Losey & Vaughan, 2006), a minority of taxa that we consider pests (*∼*1%) have an overwhelming influence on the development of population management in theory and in practice. Among the various environmentally friendly approaches that have been envisaged to control unwanted species, we focus on a research avenue that proposes the exploitation of Allee effects, i.e., the decrease in survival or reproduction at small population sizes and the consequent reduction in population growth (Dennis, 1989; Courchamp *et al.*, 1999; Stephens *et al.*, 1999; Berec *et al.*, 2007; Liebhold & Tobin, 2008). The central ideas surrounding this work are twofold: management tactics could be combined in order to (1) reduce a population size down below the Allee threshold – the population size at which the *per capita* growth rate decreases (in the case of “weak” Allee effects) or becomes negative (in the case of “strong” Allee effects) – which, in turn, increases the probability of stochastic extinction, and/or (2) amplify the mechanisms underpinning a pre-existing Allee effect to increase the Allee threshold itself (Tobin *et al.*, 2011). Capitalizing on Allee effects to manage undesirable species is particularly advantageous because it drives populations into extinction vortices without needing to eliminate every last individual.

Control methods centered on manipulating mating success as an alternative to chemical pesticides have long been recognized as desirable (e.g. Knipling, 1955; Baumhover *et al.*, 1955). The application of such control methods has been considered for managing many insect pests including the Oriental fruit fly (Steiner *et al.*, 1970), Indian meal moth and almond moths (Sower & Whitmer, 1977), and Gypsy moths (Beroza & Knipling, 1972; Knipling, 1970). Importantly, mating disruption has been successfully used to control populations with pre-existing Allee effects. The Gypsy moth (*Lymantria dispar*), for example, is one of the few insect species for which both a component (mate-finding) and demographic Allee effect have been explicitly identified (Tobin *et al.*, 2013, 2007; Johnson *et al.*, 2006). The Gypsy moth is an invasive forest pest in North America and triggered a major containment program to slow the spread toward the western United States (Sharov *et al.*, 2002a; Liebhold *et al.*, 1992). Mating disruption has been a major tactic used to control newly established low-density populations along the invasion front, with evidence supporting that it is more efficient as well as economically cheaper than classic treatments with the pesticide *Bacillus thuringiensis* (Sharov *et al.*, 2002a,b). This highlights the potential benefits of identifying other pest species that have pre-existing Allee effects and determining whether environmentally desirable forms of control may similarly be effective.

Several recent theoretical developments have focused on taking advantage of Allee effects to promote pest eradication (e.g. Boukal & Berec, 2009; Liebhold & Bascompte, 2003; Blackwood *et al.*, 2012; Yamanaka & Liebhold, 2009). These models capture the underlying population dynamics of a pest and evaluate the success of population management tactics such as culling, release of sterile males, and mating disruption to determine whether these methods can create or enhance pre-existing Allee effects (Fauvergue, 2013 provides a comprehensive review). While there is evidence that such population management strategies will be successful for populations with pre-existing Allee effects, the range of species that might benefit from these tactics may be much greater than currently known. In a meta-analysis focused on the presence of Allee effects in natural animal populations (Kramer *et al.*, 2009), terrestrial arthropods were found associated with the largest number of studies (22) and the highest proportion (77%) exhibiting an Allee effect. Mating failure at low density appeared as the most frequent mechanism. Additionally, Fauvergue (2013) found evidence supporting the presence of mate-finding Allee effects in 19 out of 34 published studies that investigated the interplay of population size and mating success in insects. Indirectly, the central role of Allee effects in insect population dynamics is supported by the efficiency of eradication programs based on the disruption of reproduction. Pest management based on the reduction of mating success via mass trapping, mating disruption with sex pheromones, or the release of sterile males has indeed proved successful in several instances (Knipling, 1955; Baumhover *et al.*, 1955; Suckling *et al.*, 2012, 2014; Krafsur, 1998).

In this article, we investigate *Wolbachia*-induced cytoplasmic incompatibility (CI) as a novel method for triggering reproductive failures and consequently bringing a pest population below its Allee threshold. *Wolbachia* are endosymbiotic bacteria that infect at least 20% of all insect species and up to two thirds in some estimations (Hilgenboecker *et al.*, 2008). *Wolbachia* have various effects on their insect hosts, the most widespread and prominent being cytoplasmic incompatibility (Stouthamer *et al.*, 1999). Under CI, matings between an infected male and a female that is either uninfected or infected with an incompatible cytotype result in offspring mortality during embryonic development. Fitness advantages of infected females as well as maternal inheritance are key features that promote invasion of *Wolbachia* into a host population: above a threshold frequency, a given *Wolbachia* strain is expected to invade until near-fixation (Barton & Turelli, 2011; Hancock *et al.*, 2011; Caspari & Watson, 1959; Hoffmann & Turelli, 1997; Turelli & Hoffmann, 1991). As a result of the associated CI and subsequent reduction in reproductive rate, *Wolbachia* invasion via the release of infected hosts is a candidate biological control agent against arthropod pests (Bourtzis, 2008).

In practice, there are multiple ways to implement a management strategy centered on inducing CIs via introduction of *Wolbachia*. For example, similar to the use of “Sterile Insect Technique” (SIT), males bearing a *Wolbachia* strain incompatible with that of the target population can be released in large numbers. CIs arising from the mating of females and infected males would substantially limit the total offspring in the subsequent generation, resulting in a decrease in overall population growth rate and thereby increasing the possibility of local population extinction (Laven, 1967; Zabalou *et al.*, 2004; Atyame *et al.*, 2015). Incompatible males can be obtained via transfection, even between completely different species of host insects (e.g. Braig *et al.*, 1994; Hoffmann *et al.*, 2011). At the population level, the conceptual underpinnings for mass-releases of incompatible males do not depart from that of SIT, for which interactions with the Allee effect have already been thoroughly analyzed (Boukal & Berec, 2009; Yamanaka & Liebhold, 2009; Fauvergue, 2013; Barclay & Mackauer, 1980; Barclay, 1982; Berec *et al.*, 2016; Lewis & Van Den Driessche, 1993).

An alternative management tactic using CI relies on the inoculation of a relatively small number of insects of both sexes with a *Wolbachia* strain incompatible with that of the target population. This method is investigated in the model introduced in Dobson *et al.* (2002), which combines insect population dynamics with releases of individuals infected with*Wolbachia*. During a successful invasion of *Wolbachia*, a transient reduction in the insect population size occurs. This decline results from the temporary increase in the fraction of incompatible matings, which peaks in the midst of the invasion process. Therefore, systematic introductions of different *Wolbachia* cytotypes could be applied to artificially sustain an unstable coexistence of multiple incompatible infections within an insect population, allowing the population size to be reduced and maintained at low levels (Dobson *et al.*, 2002).

Our goal is to determine when the latter implementation of *Wolbachia* introductions can drive a population to extinction in the presence of Allee effects. Specifically, we derive a mathematical model built upon Dobson *et al.*’s (2002) approach of CI management that additionally accounts for Allee effects as well as environmental and demographic stochasticity. We also consider mating disruption in our model as a potential complementary tactic. We use this model to address three primary questions: (1) What is the influence of Allee effects present within a host population on *Wolbachia* invasion dynamics? (2) What is the influence of cytoplasmic incompatibility on the demographic Allee effect? (3) What is the influence of a combination of *Wolbachia*-induced CI, Allee effects, mating disruption, and stochasticity on the probability of host extinction?

## Methods

### Population model

Our model extends the framework introduced by Dobson *et al.* (2002) by accounting for pre-existing Allee effects, the release of pheromone sources as a method of mating disruption, as well as both demographic and environmental stochasticity. In this section, we first introduce a model that considers the population dynamics in the absence of individuals infected with *Wolbachia*.

We considered populations such that the dynamics can be modeled in discrete time with non-overlapping generations. The population model explicitly tracks the total population size at each time *t*, given by *N_t_*. Our population model can be expressed in terms of either census size or density and hereafter, we refer to our model in terms of size. Therefore, while the deterministic model can take non-integer values, population sizes less than one are considered extinct. In contrast, the stochastic model forces integer population sizes. We assume that each time step can be broken into two stages: the first (at time *t*+0.5) captures reproduction, and the second (at time *t* + 1) captures density dependent survivorship of offspring to adults. The total number of offspring is given by:

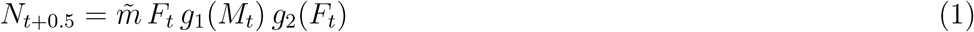

where *m*̃ is the maximum *per capita* female fecundity, *F_t_* is the number of females in the population, and *M_t_* is the number of males. *g*_1_(*M_t_*) captures a component Allee effect that results from the failure of mates finding one another at low densities such that:

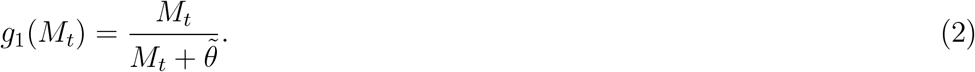

Here, *θ*^̃^ measures the strength of the Allee effect; a convenient interpretation of this term is that *θ*^̃^ represents the number of males at which half of the females successfully find a mate or, equivalently, the maximum mating rate is reduced by half (Boukal & Berec, 2009).

The function *g*_2_(*F_t_*) in Eqn. 1 captures the decline in fecundity resulting from techniques to control populations via mating disruption. We assume that pheromone sources are introduced into the environment and maintained in the population at a fixed number *P*^̃^ and only a fraction *F_t_*/(*F_t_* + *P*^̃^) males successfully find a mate (Fauvergue, 2013), or:

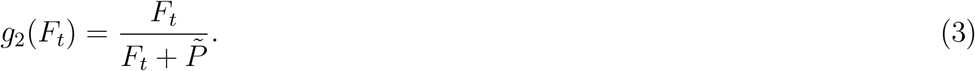

We now assume that there is a 50:50 sex ratio (i.e. *F_t_* = *M_t_* = *N_t_*/2) and define constants *m* = *m*̃/2 (which is now the overall *per capita* fecundity), *θ* = 2*θ*^̃^, and *P* = 2*P*^̃^. Now, we can write both *g*_1_ and *g*_2_ as functions of *N_t_* and Eqn. 1 is now:

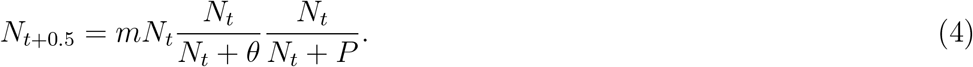

For a simpler biological interpretation, we consider *θ* relative to the carrying capacity of the population in the absence of any control (denoted as *K*) as an indicator of the intensity of the Allee effect (i.e. *θ/K*). Now, when *θ* = *K* (or, equivalently, *θ/K* = 1) half of the *total* population successfully mates at its carrying capacity. Hereafter, we refer to *θ/K* as the “relative strength of the Allee effect” and restrict its values to the range (0 – 0.5), with 0.5 likely on the upper end of biologically reasonable values for *θ/K*. Similarly, we consider *P*, the number of pheromone sources, relative to *K*.

Finally, we assume that survivorship of offspring to adults is density dependent so that:

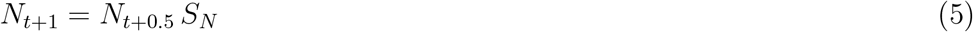

where

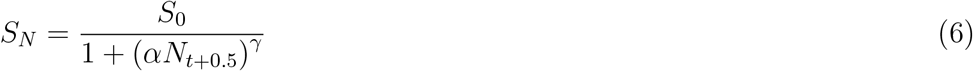

and the constant *α* is related to the carrying capacity (more details provided in the Supplementary Information S1), *γ* is related to intraspecific competition, and *S*_0_ is survivorship in the absence of intraspecific competition (Slatkin & Smith, 1979). Combining equations (4)-(7), we are left with the model:

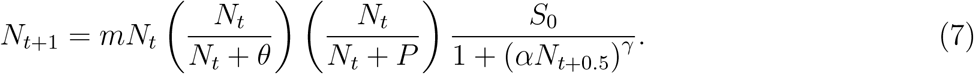

Our goal is to consider the dynamical consequences of population control methods (i.e. *Wolbachia* introductions and the release of pheromone sources). Therefore, we analytically determine the equilibrium values for the population model (7) in the absence of these management tactics (*P* = 0). Throughout the remainder of the paper, we distinguish between the “maximum reproductive rate” (which is given by *mS*_0_ and therefore accounts for both fecundity and density independent survivorship) and the overall “reproductive rate” (which is the reproductive rate in the presence of all other demographic processes in our model, namely the Allee effect). As detailed in the Supplementary Information (S1), we find three equilibria (assuming the maximum reproductive rate is greater than one and setting *γ* = 1). The first equilibrium corresponds to population extinction, and the second two are given by:

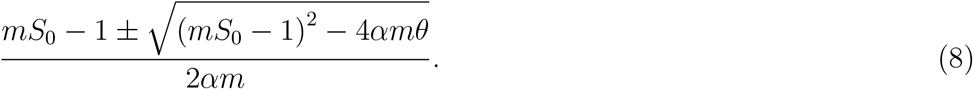

When Allee effects are not present (*θ* = 0) these equilibria collapse to a single equilibrium (the carrying capacity *K*). In contrast, for sufficiently large Allee effects these equilibria no longer exist and the reproductive rate is always less than one; consequently, the population will be driven to extinction independent of its initial size. However, when the value of *θ* is between these extremes, the smaller equilibrium corresponds to the Allee threshold and the larger equilibrium corresponds to the carrying capacity. The structure of these equilibria are integral to the insect species that we are considering: there is a carrying capacity and a strong Allee effect for positive values of *θ* up to a threshold. We therefore conjecture that an alternative form of density dependence that captures these properties will yield qualitatively similar results.

Based on this deterministic framework, we build in environmental and demographic stochasticity. We assume that environmental stochasticity leads to variation in the population’s fecundity between generations (Melbourne & Hastings, 2008). Therefore, we adapt the methods of Schoener *et al.* (2003) and account for environmental stochasticity by rewriting Eqn. 4 as:

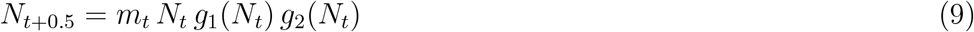

where the fecundity *m_t_* is drawn at each generation from a normal distribution with mean *m* (that is truncated so that *m* ≥ 0) and variance (*σ*^2^) that scales with the mean. In the main text, we fix the variance so that it is equal to the mean; however, a sensitivity analysis of the magnitude of the variance is provided in the Supplementary Information (S3). Further, we note that gamma-distributed environmental stochasticity is a commonly used alternative choice; under our parameterization, the probability density function (pdf) is nearly identical for both normal- and gamma-distributed stochasticity and would therefore yield similar results.

Demographic stochasticity results from variation in fecundity at the individual level (Melbourne & Hastings, 2008). In the absence of demographic stochasticity, the total number of individuals that successfully reproduce is given by:

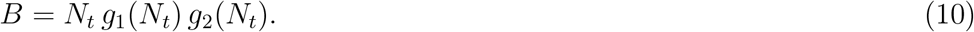

We assume that each of these individuals at a given time *t* reproduces with fecundity *m_t_* (as described above), and the total number of eggs produced is a Poisson random variable (Melbourne & Hastings, 2008). Since the sum of independent Poisson random variables is also a Poisson random variable, the total offspring of all adults is:

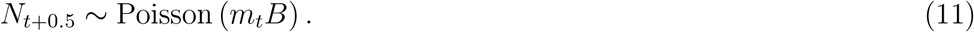

Finally, we include stochasticity in density dependent survivorship, again following Melbourne & Hastings (2008). Given that *S_N_* (as defined in Eqn. 7) is the probability that offspring survive to adults, we assume that survivorship is binomially distributed so that:

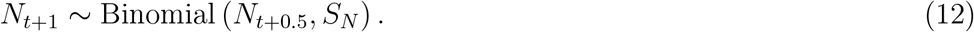

### Infection dynamics

We consider the infection dynamics of up to two different cytotypes of *Wolbachia* (referred to as cytotypes *X* and *Y*) and denote the number of uninfected individuals as *W*. Note that all variables and parameters with subscripts *X* (or *Y*) are related to cytotype *X* (or *Y*). This model is adapted from Dobson *et al.* (2002); therefore, we use similar notation throughout.

In the presence of a single cytotype of *Wolbachia*, there are only unidirectional cytoplasmic incompatibilities (CI); in contrast, in the presence of multiple cytotypes there may be bi-directional CI. We first introduce the case of a single cytotype and then extend the model to include two cytotypes. Below we describe the mathematical formulation of the infection dynamics, and Table 2 summarizes these processes.

**Table 1:** List of model parameters. All figures use these parameter values, unless otherwise stated. Parameter values with a “ ” have associated sensitivity analyses (as discussed in the main text) in the Supplementary Information (S3). [1] refers to the reference Dobson *et al.* 2002, and [2] refers to Charlat *et al.* (2005).

| Parameter | Description | Value | Source |
| --- | --- | --- | --- |
| $N_0$ | initial population size | varies 1–$K$ | |
| $m, m_t$ | *per capita* fecundity (realized, expected) | 25 | |
| $\sigma^*$ | standard deviation of fecundity | 5 | |
| $S_0^*$ | survivorship in absence of competition | varies 0.08–0.2 | |
| $\alpha$ | related to carrying capacity | 0.00002 | [1] |
| $\gamma$ | related to intraspecific competition | 1 | [1] |
| $\mu_X^*, \mu_Y$ | maternal transmission failure | 0.03 | [1] |
| $F_X^*, F_Y$ | relative fecundity of infected individuals | 0.95 | [1] |
| $H_X, H_Y$ | proportion of offspring surviving CI | 0.05 | [2] |
| $\theta$ | strength of Allee effect | varies 1–0.5$K$ | |
| $P$ | number of pheromone sources | varies 0–0.15$K$ | |

**Table 2:**
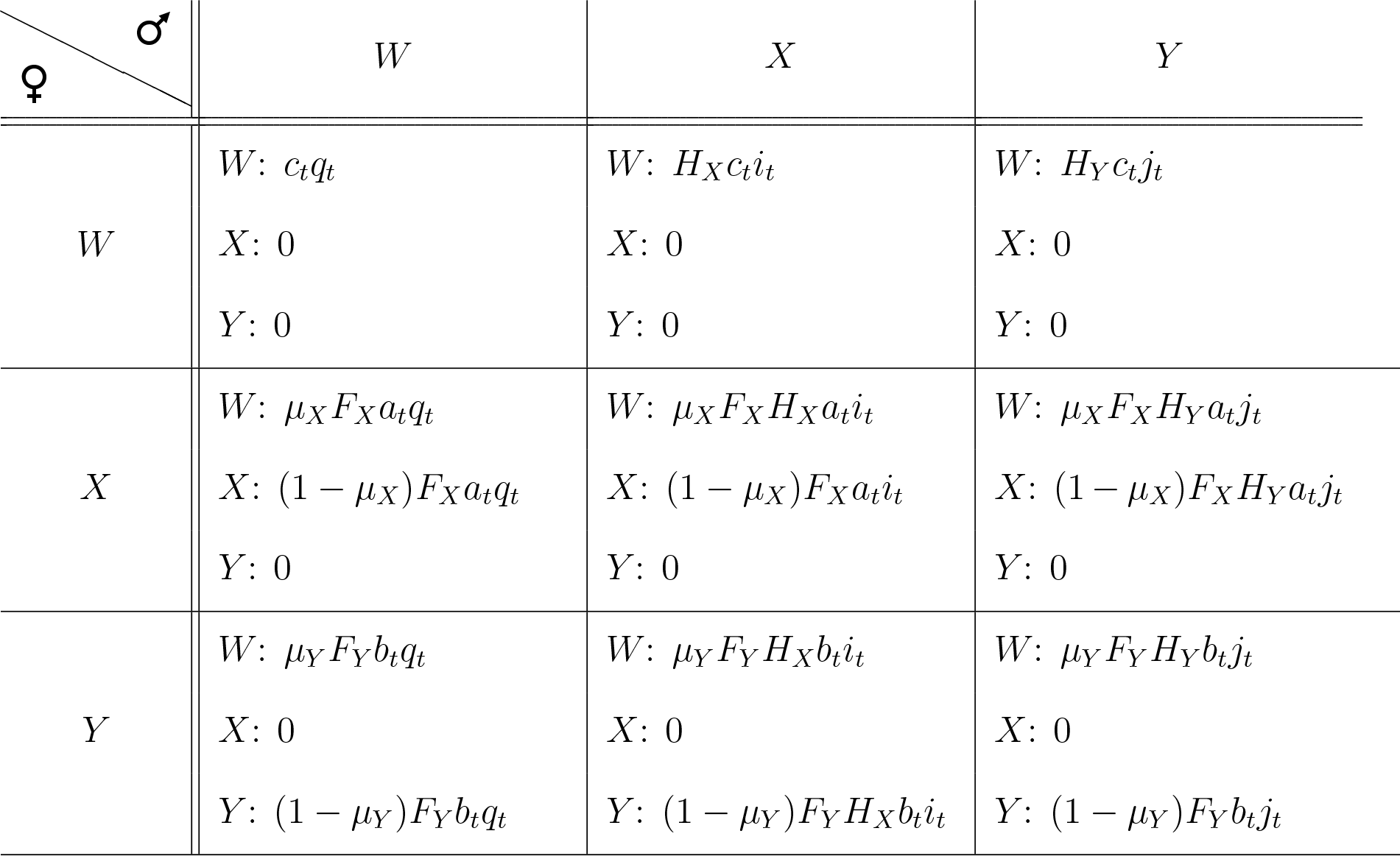
Summary of *Wolbachia* transmission and its effects on reproduction in its host population. The first column states the maternal *Wolbachia* cytotype and the first row states the paternal *Wolbachia* cytotype. Each box in the table corresponding to a particular pairing between a female and male provides that proportion of the offspring from that pair that are uninfected (*W*), infected with cytotype *X*, and infected with cytotype *Y*.

#### One cytotype

To capture the *Wolbachia* dynamics, we first determine the proportions of infected and uninfected individuals in the population at time *t*. For example, if there are *W_t_* uninfected individuals and *X_t_* infected with cytotype *X* then, under the assumption that there is a 50:50 sex ratio, the fraction of females infected with cytotype *X* at *t* + 0.5 is given by:

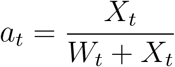

Similarly, we find the fraction *i_t_* of all males that are infected (where *i_t_* = *a_t_*), the fraction *q_t_* of all males that are uninfected, and the fraction *c_t_* of all females that are uninfected (again *q_t_* = *c_t_*).

Based on the proportions of uninfected and infected individuals in the population, we can now determine the fraction of offspring that are infected. Crosses between pairs with an infected female suffer a fecundity loss due to infection (1*−F_X_*), where *F_X_* is the relative fecundity of infected individuals. Vertical transmission of *Wolbachia* occurs maternally and we assume that transmission is successful with probability (1 *− µ_X_*), where *µ_X_* is the probability of transmission failure. In the instance of *Wolbachia*-induced CIs, crosses between infected females and uninfected males in addition to crosses between infected males and infected females give rise to infected offspring. The proportion of viable offspring that are infected with cytotype *X* after reproduction (i.e. at time *t* + 0.5) is therefore given by:

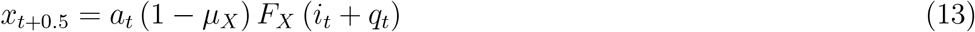

where a lowercase *x* is used to denote proportion rather than number. Second, we can identify the proportion of viable offspring that are uninfected (*w_t_*_+0.5_). Uninfected individuals can arise from crosses between both uninfected females and uninfected males. Further, matings between both infected females and infected males can yield viable uninfected offspring. This results from failure to vertically transmit *Wolbachia* to their offspring. When one type of *Wolbachia* is present within a population, then only unidirectional cytoplasmic incompatibility (CI) is possible. This type of CI occurs through matings between infected males and uninfected females. Therefore, we assume that all but a fraction *H_X_* of pairings between infected males and uninfected females undergo CI. Additionally, offspring from pairings between infected males and infected females that fail to transmit *Wolbachia* are subject to CI-induced mortality. The proportion of viable offspring that are not infected with *Wolbachia* following reproduction is therefore given by:

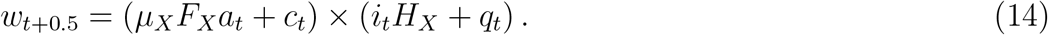

Notice that due to cytoplasmic incompatibilities and the fecundity cost due to infection with *Wolbachia*, the fraction of the offspring that are viable (*x_t_*_+0.5_ + *w_t_*_+0.5_) is less than one. Therefore, the total number of offspring as governed by Eqn. 4 can be rewritten as:

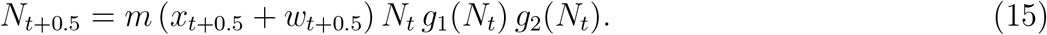

In other words, the product *g*_1_(*N_t_*)*g*_2_(*N_t_*) captures the total fraction of adults at time *t* who successfully find a mate, and the sum *x_t_*_+0.5_ + *w_t_*_+0.5_ is the fraction of all offspring that are viable. Finally, as described in the previous section, density dependent mortality limits the total number of adults at time *t* + 1 (Eqn. 7).

#### Two cytotypes

In addition to releasing a single cytotype of *Wolbachia*, we consider a scenario in which a second cytotype is introduced. When two cytotypes of *Wolbachia* are present within a population, bidirectional CI occurs when a male with one cytotype mates with a female infected with an incompatible *Wolbachia* cytotype. Similar to the previous section, we assume that a fraction *H_X_* (or *H_Y_*, depending on the infection type of the male and female) of offspring survives. Offspring of pairings between infected males and females of either cytotype that fail to transmit are again subject to CI-induced mortality.

Therefore, in the presence of two strains we rewrite Eqn. 13 as:

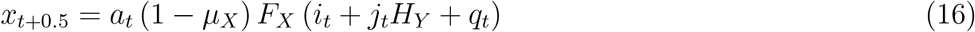

where *j_t_* is the fraction of males infected with cytotype *Y*. Similarly, the proportion of viable offspring infected with cytotype *Y* following reproduction is given by:

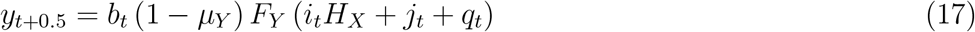

where *b_t_* is the fraction of females infected with cytotype *Y*. The proportion of viable uninfected offspring is now given by:

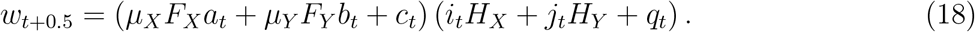

### Parameterization

Here, we define the maximum reproductive rate as the product of the *per capita* fecundity and density independent survivorship (i.e. *mS*_0_). In other words, this is the reproductive rate in the absence of Allee effects and in the limit as the population size approaches zero. Our parameterization of the population model is based on both the parameterization used in Dobson *et al.* (2002) and common ranges for insect populations (see Table 1). For example, Hassell *et al.* (1976) cites finite net rates of increase (defined as fecundity discounted by density independent mortality and therefore equivalent to *mS*_0_ here) ranging from 1.3 − −75 (with 22 of the 24 species investigated in the range 1.3 − −13.5) which is consistent with our parameterization (e.g. see Fig. 2 caption and the associated results as *mS*_0_ is varied). Additionally, several parameters vary for our analysis including the strength of the Allee effect, *θ*, and the initial population size. However, our results are intended to assess the general qualitative behavior of *Wolbachia* introductions and consequently the actual implementation of such management tactics would require a detailed analysis and parameterization specific to the target population and cytotype.

Our parameterization for the infection dynamics is based on values that are reasonable for *Wolbachia* cytotypes (Hoffmann & Turelli, 1997; Dobson *et al.* (2002); Charlat *et al.*, 2005). In the main text, we assume that fecundity loss, transmission failure, and survival of CI are equal between all cytotypes. However, the Supplementary Information (S3) provides an analysis of the dynamics when the introduced cytotypes are not identical. S3 additionally provides sensitivity analyses for parameters related to the population model, including the strength of environmental stochasticity and the maximum reproductive rate.

## Results

In the following sections we first test our model against well-established results related to *Wolbachia* invasion as a method of model validation, establish the relationship between *Wolbachia* and the strength of the Allee effect, and finally evaluate the potential for the release of infected insects to control a population.

### Model validation

We first determine whether our model captures the same features of the important earlier work (Hoffmann *et al.*, 1990; Turelli & Hoffmann, 1991; Hancock *et al.*, 2011). Hoffmann *et al.* (1990) derived an analytic expression for the expected equilibrium infection frequencies. After adjusting their notation to match ours and simplifying, the equilibrium infection frequency (which we denote as *p*) for a single cytotype of *Wolbachia X* should satisfy the equation:

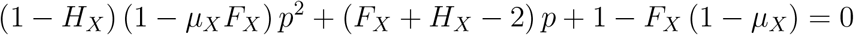

Their work predicts that there is an unstable equilibrium referred to as the *Wolbachia* invasion threshold; cytotypes with initial infection frequencies above this threshold will increase until reaching the higher stable equilibrium, indicating a successful invasion. Following Charlat *et al.* (2005), we considered invasion dynamics by estimating the infection frequency at generation *t* + 1 as a function of the frequency at generation *t*. Doing so allows us to create a simple graphical representation of the stable and unstable equilibria (Fig. 1). Unless stated otherwise, the default parameter values are listed in Table 1.

**Figure 1.**
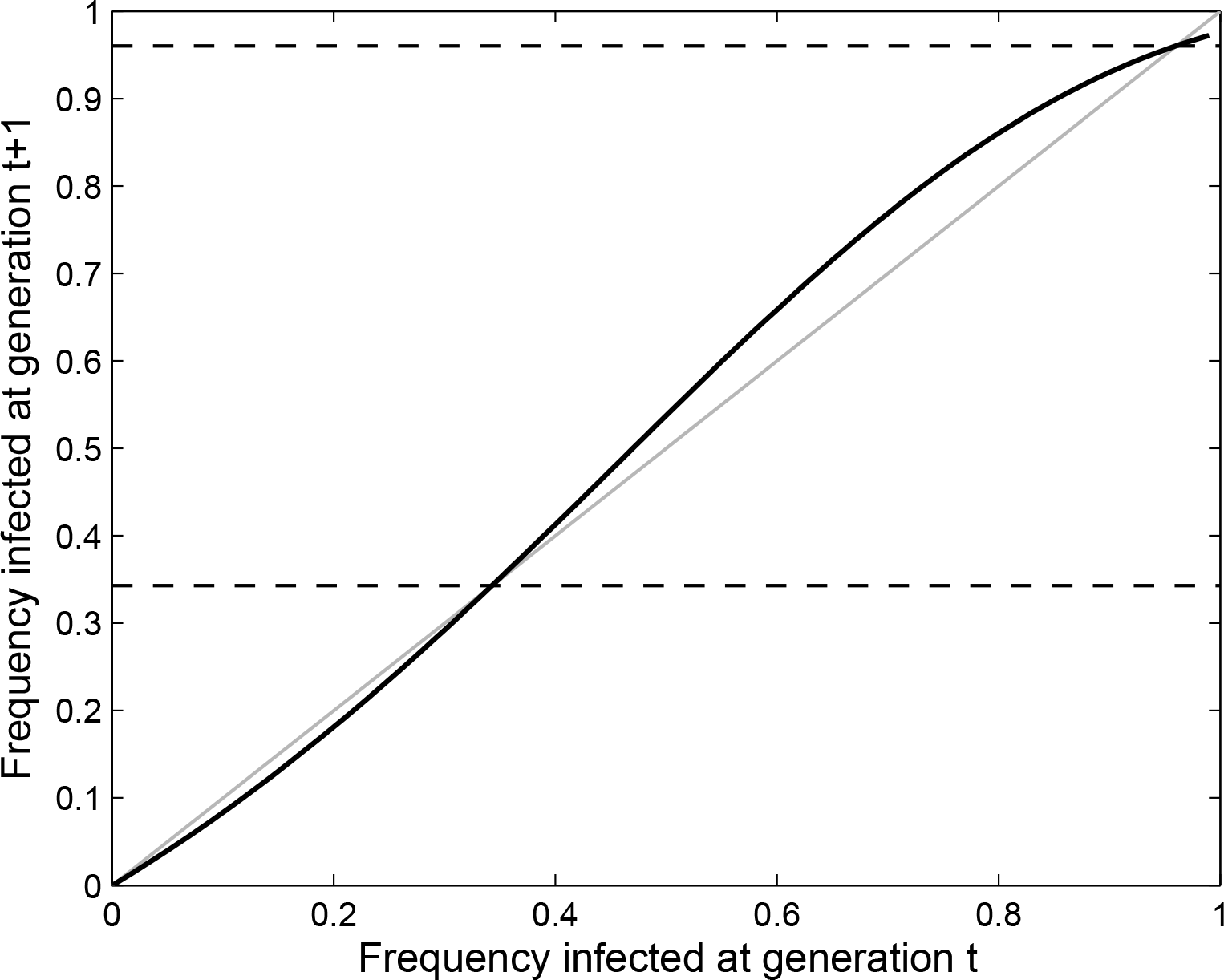
Verification that our model accurately predicts the invasion threshold as analytically determined in Hoffman *et al.* 1990. Here, we ignore Allee effects, stochasticity, and assume *P* = 0. The thick black curve is the frequency of infected individuals at time *t* + 1 given the frequency at *t*; equilibria occur when this curve and the gray line (which corresponds to the case that the frequency at generations *t* and *t* + 1 are equal) intersect. When the black curve lies above the gray line, the infection frequency is increasing; similarly, the infection frequency is decreasing when the black curve falls below the gray line. The dashed lines indicate the analytically predicted equilibrium. The smaller intersection is an unstable equilibrium that defines the invasion threshold: individuals introduced at a frequency higher than this threshold will successfully invade the population and approach the higher stable equilibrium. In this figure, we use more extreme values for parameters related to CI to more clearly demonstrate the location of the invasion threshold (specifically, *µ_X_* = 0.2, *H_X_* = 0.1). Here, *S*_0_ = 0.15.

Our simulation results are consistent with the analytically derived equilibrium infection frequencies (Fig. 1). This verifies that our simulations are in line with the behavior that we would like to capture from our model and are consistent with the results observed in Charlat *et al.* (2005). This is not surprising given that our model makes similar assumptions on the mechanisms driving *Wolbachia* invasion dynamics (e.g. fecundity loss and cytoplasmic incompatibilities). In contrast to earlier studies, our population model is dynamically different because of the inclusion of Allee effects and pheromones. Therefore, we determined the relationship between the invasion threshold and these features of the model. We found that the *Wolbachia* invasion threshold is not affected by Allee effects nor by the application of pheromones to the host insect (not shown). This is important to note because in all of our simulations and analyses, the invasion threshold does not vary as *θ* and *P* are adjusted. Finally, we note that the invasion threshold is not affected by any of our demographic parameters (i.e. *m*, *α*, *γ*, and *S*_0_; not shown.)

### The effect of *Wolbachia* on the Allee threshold

To determine the dynamical effects of the presence of *Wolbachia* infection within a population, we find the Allee threshold in insect populations both in the presence and absence of infection. In this section, we ignore stochasticity as well as the release of pheromones (*P* = 0). For a given proportion of infected individuals, we calculate the reproductive rate between two consecutive generations (i.e. *N_t_*_+1_*/N_t_*) across all population sizes (Fig. 2). The equilibria for our population model occur when *N_t_*_+1_ = *N_t_*, and there are three equilibria for the parameters used to produce this figure: the first corresponds to population extinction (stable), the second is the Allee threshold (unstable), and finally the third is the carrying capacity (stable).

**Figure 2.**
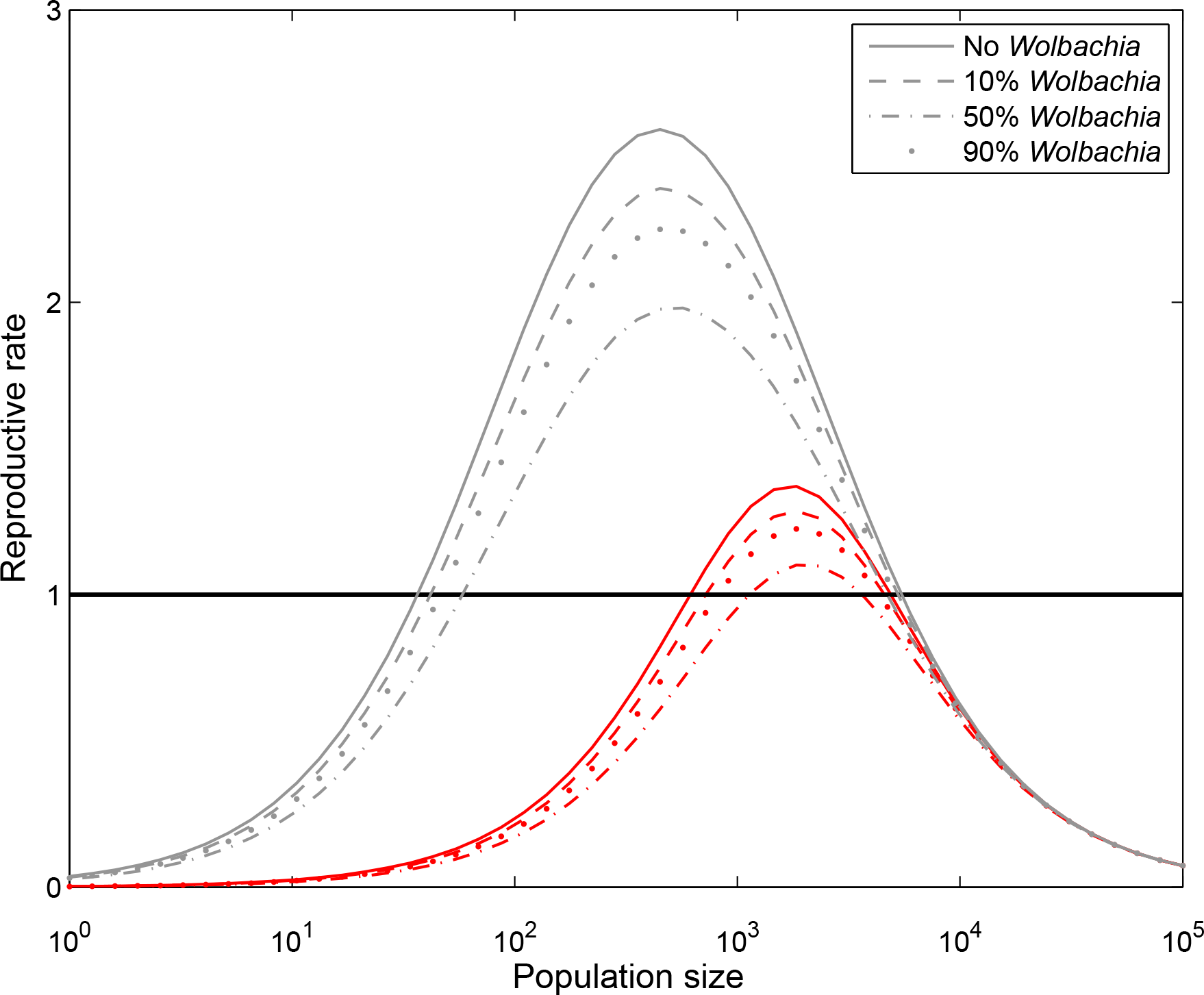
Reproductive rate as a function of population size when *θ* = 100 (gray) and *θ* = 1500 (red). Values above one correspond to population growth, and values below one correspond to decline. The populations corresponding to the solid lines have no *Wolbachia*-infected individuals, populations with dashed lines have 10% of the population infected, dash-dotted lines have 50% of the population infected, the population infected, dash-dotted lines have 50% of the population i0.15.

In addition to considering the population model in the absence of *Wolbachia*-infected individuals, we calculated the reproductive rates when the population is comprised of 10%, 50%, and 90% infected individuals (Fig. 2). Given our parameterization, the frequency of infected individuals is chosen to lie above the invasion threshold (which is *∼* 8.5%); therefore, this figure captures the dynamics between two consecutive generations during the replacement process when the population contains the specified distribution of infected and uninfected individuals. As a consequence of cytoplasmic incompatibilities, there is an increase in the Allee threshold. Additionally, there is a slight decrease in the carrying capacity that results from the fecundity loss associated with *Wolbachia* infection. Consequently, the maximum reproductive rate decreases as the proportion of infected individuals increases. Finally, the changes in reproductive rate are most significant during the replacement process (e.g. when 50% is infected) and they diminish when the infection is close to fixation (e.g. 90%). The proportion of *Wolbachia*-infected individuals has a significantly smaller effect on the location of the Allee threshold than the strength of the mate finding Allee effect itself (Fig. 2).

### Implications for population management

#### Deterministic results

In this section, we characterize implications for population management through the release of *Wolbachia*-infected individuals into an insect population. As observed by Dobson *et al.* (2002), there should be a transient decline in the population size during the replacement of uninfected hosts by *Wolbachia*-infected individuals. Therefore, we find the magnitude of this decline in the presence of Allee effects to determine the conditions under which the replacement process brings the population size below the Allee threshold in a deterministic setting, thereby forcing extinction. This is achieved by running our model over a range of values for the strength of the mate-finding Allee effect (*θ*). We assume that the initial population size is at its carrying capacity (which is found analytically, as shown in the Supplementary Information S1).

We find the minimum population size (relative to *K*) over 50 generations following the introduction of one cytotype (Fig. 3B) and two cytotypes (Fig. 3C). Here, values of zero for the minimal population size indicate that the transient reduction in population size brought the population below the Allee threshold, therefore leading to deterministic extinction. The first cytotype is always released in the second generation, and the release of the second cytotype is optimized so that the release occurs in the generation that causes the largest decline in population size (see Supplementary Information S2 for a detailed explanation of the optimization scheme). While in the main text we assume that all cytotypes have the same infection properties, this assumption is challenged in the Supplementary Information (S3) and our qualitative results are unchanged. To ensure that the introduction size is above the invasion threshold, in all simulations we assume that the introduction is large enough so that the initial infection frequency is 10%. This value lies just above the actual threshold of *∼* 8.5% associated with our parameter values. Therefore, the number of infected individuals introduced in our simulations directly depends on the current host population size.

**Figure 3.**
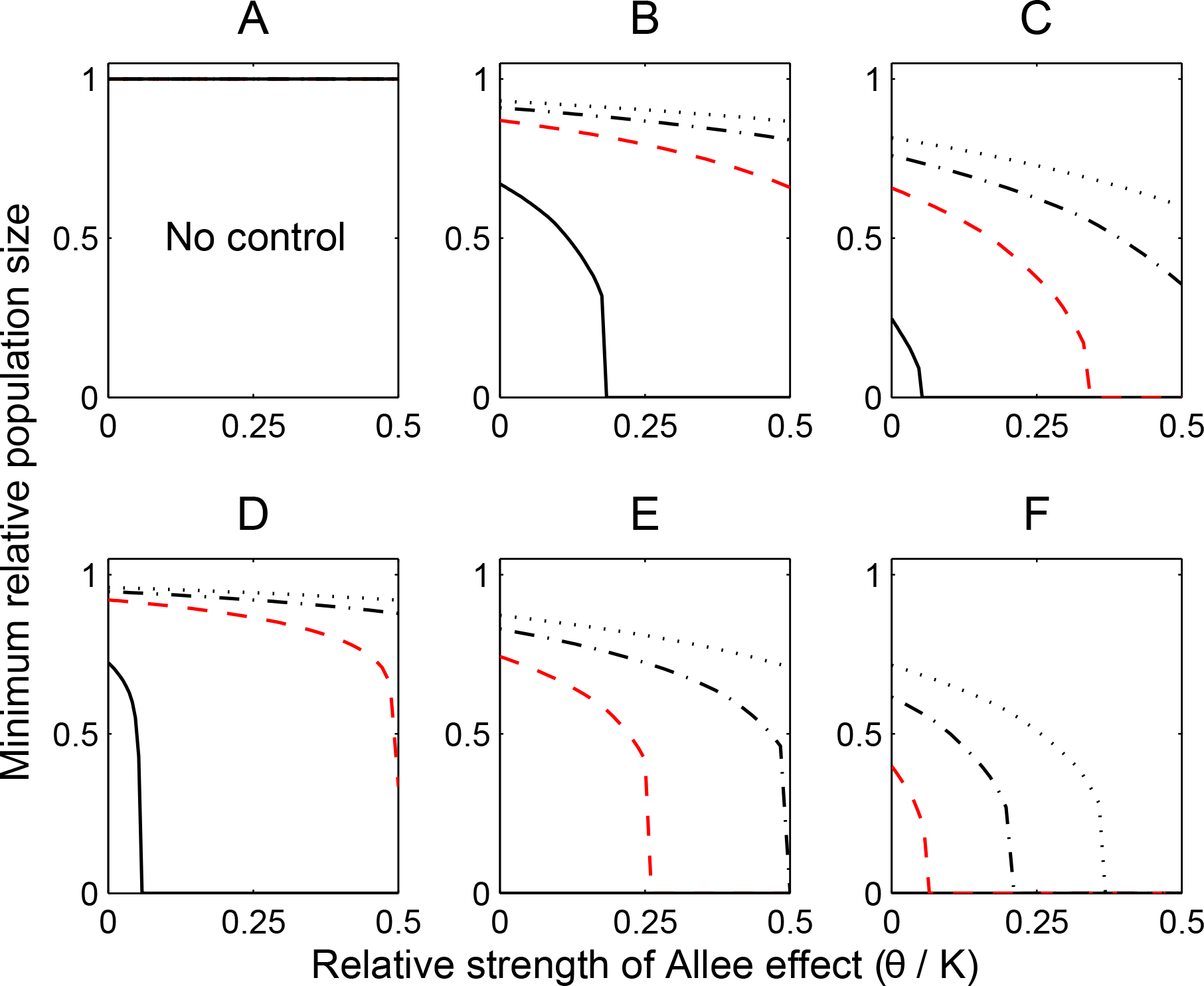
Deterministic results. (A) no control (so the population size remains at *K*); (B) single *Wolbachia* introduction; (C) two introductions. Plot displays the minimum population size relative to *K* over 50 generations assuming that *N*_0_ = *K* versus the relative strength of the Allee effect (*θ/K*). The solid line has *S*_0_ = 0.08 (maximum reproductive rate of 2 in the absence of Allee effects as in Dobson *et. al*, 2002), the dashed red line has *S*_0_ = 0.15 (maximum reproductive rate of 3.75 in the absence of Allee effects), dash-dotted line has *S*_0_ = 0.2 (maximum reproductive rate of 5 in the absence of Allee effects), and dotted line has *S*_0_ = 0.25 (maximum reproductive rate of 6.25 in absence of Allee effects). (D)-(E) are identical to (A)-(C), respectively, but it is additionally assumed that pheromone sources are held at a fixed level such that *P/K* = 0.1. In all plots, each release is created so the initial infection frequency of that cytotype is 10%. The first release is at generation one, and the second release is determined by the optimization scheme detailed in Section S2 in the ESM. In (E)-(F), the absence of the solid line indicates that extinction is achieved for all values of *θ/K*. Curves corresponding to *S*_0_ = 0.15, the value used in all subsequent plots, are highlighted in red to further emphasize the synergy between management tactics.

To determine the success of releases under varying reproductive rates, we replicated the results for four different values of *S*_0_. For all reproductive rates, the release of two incompatible cytotypes of *Wolbachia* is substantially more effective than a release of a single cytotype. In fact, release of a single cytotype is only effective in driving population extinction when reproductive rates are small. This observation holds more generally: the effectiveness of *Wolbachia* introductions increases as the maximum reproductive rate decreases (Fig. 3B-C).

These results suggest that CI management may fail in species with high reproductive rates or alternatively in these species, complementary tactics that either decrease the population size or increase the Allee threshold may be required to amplify the effects of *Wolbachia* introductions. Therefore, we consider the use of mating disruption through the release of sex pheromones (*P*) as a supplemental management tactic (as in Eqns. (4) and (3)). We determine the minimum population size following no introduction, one introduction, and the introduction of two cytotypes while additionally assuming there is a fixed number of pheromone sources present. These results suggest that combining both methods is significantly more effective than either tactic alone (Fig. 3D-F). The associated dynamics can be visualized by considering a plot of the population size with time (Fig. 4).

**Figure 4.**
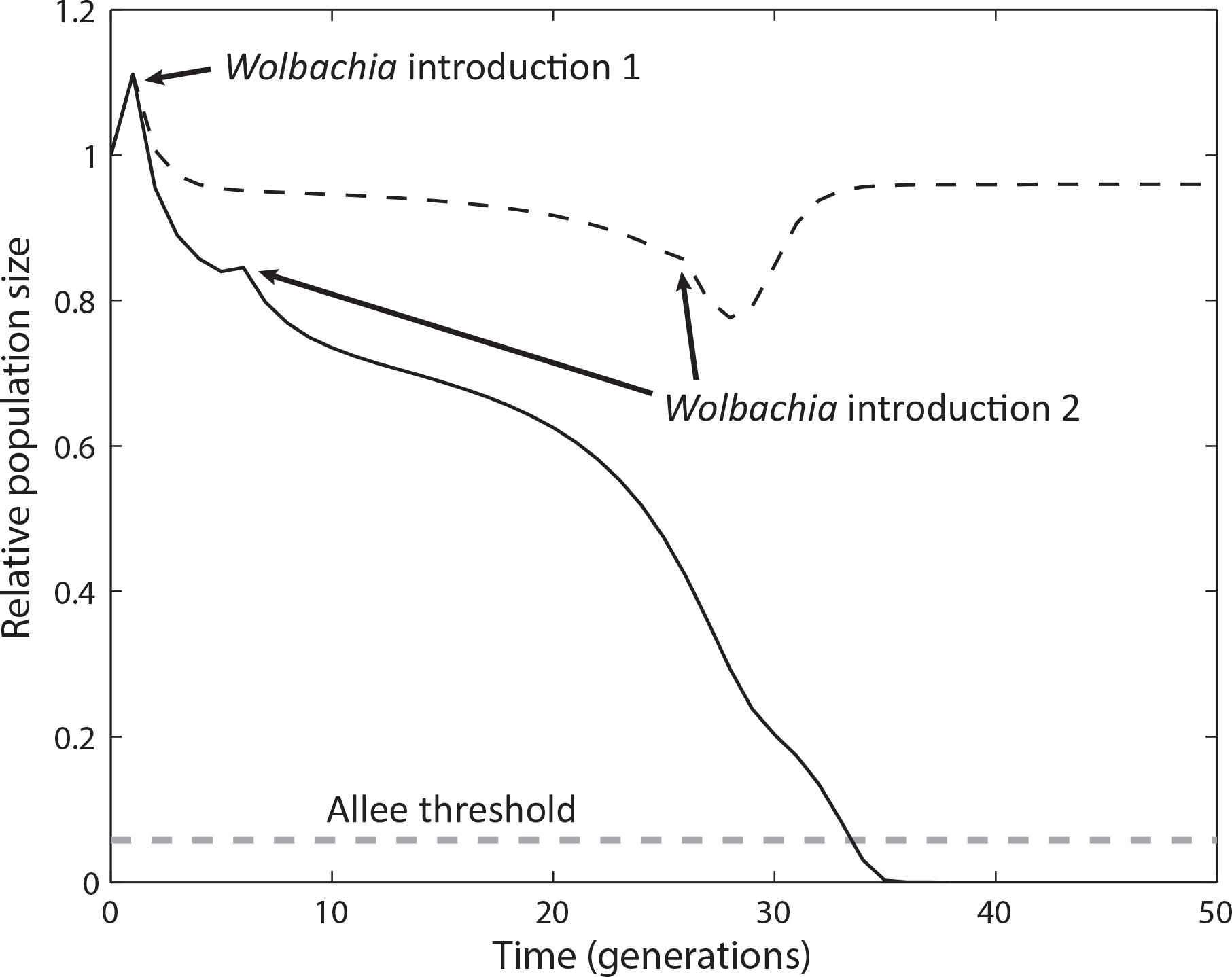
Sample trajectories of the population dynamics (relative to *K*) when two incompatible cytotypes of *Wolbachia* are introduced into a population when *S*_0_ = 0.15. The relative strength of the Allee effect is set to *θ/K* = 0.15, which corresponds to an Allee threshold of *∼*5.8% of the carrying capacity (as indicated by the horizontal dashed gray line). The black dashed line assumes that there are no pheromone sources, and the solid line assumes that the relative number of pheromone sources *P/K* is 0.1. As also evidenced in Fig. 3 (e.g. by considering the dashed curve in Fig. 3A-B when *θ/K* = 0.15), intervention with *Wolbachia* introductions only under these parameter conditions is not successful. A combination of *Wolbachia* introduction and pheromone sources, however, is successful in achieving population extinction.

#### Stochastic results

The interplay of Allee effects and stochasticity can be especially important at low population sizes, when the population is at higher risk of stochastic extinction. Therefore, in this section we determine the ability of *Wolbachia* and mating disruption to drive populations with variable initial population sizes to extinction in the presence of Allee effects and stochasticity.

To achieve this, we determine the probability of extinction based on 500 realizations of the stochastic model (i.e. Eqns. 9-12) over all relevant combinations of the initial population size and strength of the Allee effect (i.e. the initial population size is at most at 80% of carrying capacity and the relative Allee effect, *θ/K* is no greater than 0.5). To determine the relative roles of environmental and demographic stochasticity, we simulate the model while including both types of stochasticity as well as demographic stochasticity alone. Further, we find the extinction probability under three scenarios: no introduction of *Wolbachia*-infected individuals, introduction of one cytotype, and the introduction of two incompatible cytotypes. As in the deterministic setting, we assume that the introduction of the first cytotype occurs at the second generation. When two cytotypes are introduced, the generation of the second release is determined in the same way as it is found in the deterministic setting: the second introduction is optimized for each realization so that it occurs in the generation (up to 25 generations) that creates the largest transient decrease in population size resulting from the *Wolbachia* introduction. The number of generations between releases increases as the strength of the mate-finding Allee effect decreases (see Supplementary Information S2). As before, each release is implemented so that the proportion of infected individuals of a given cytotype is 10% (just above the invasion threshold).

Here, we consider populations with relatively high reproductive rates (maximum reproductive rate 3.75); the Supplementary Information (S3) provides a sensitivity analysis for lower reproductive rates. We find that the introduction of a single cytotype of *Wolbachia* leads to negligible increases in the extinction probability across all values of the relative strength of the Allee effect (Fig. 5A-B). However, environmental stochasticity increases the uncertainty associated with extinction near the Allee threshold, and there are more apparent increases in extinction probability for *Wolbacia* introductions for very strong Allee effects (Fig. 5D-E). As described in the previous section, the success of *Wolbachia* releases increases for lower reproductive rates. This finding holds in the stochastic setting (see S3).

**Figure 5.**
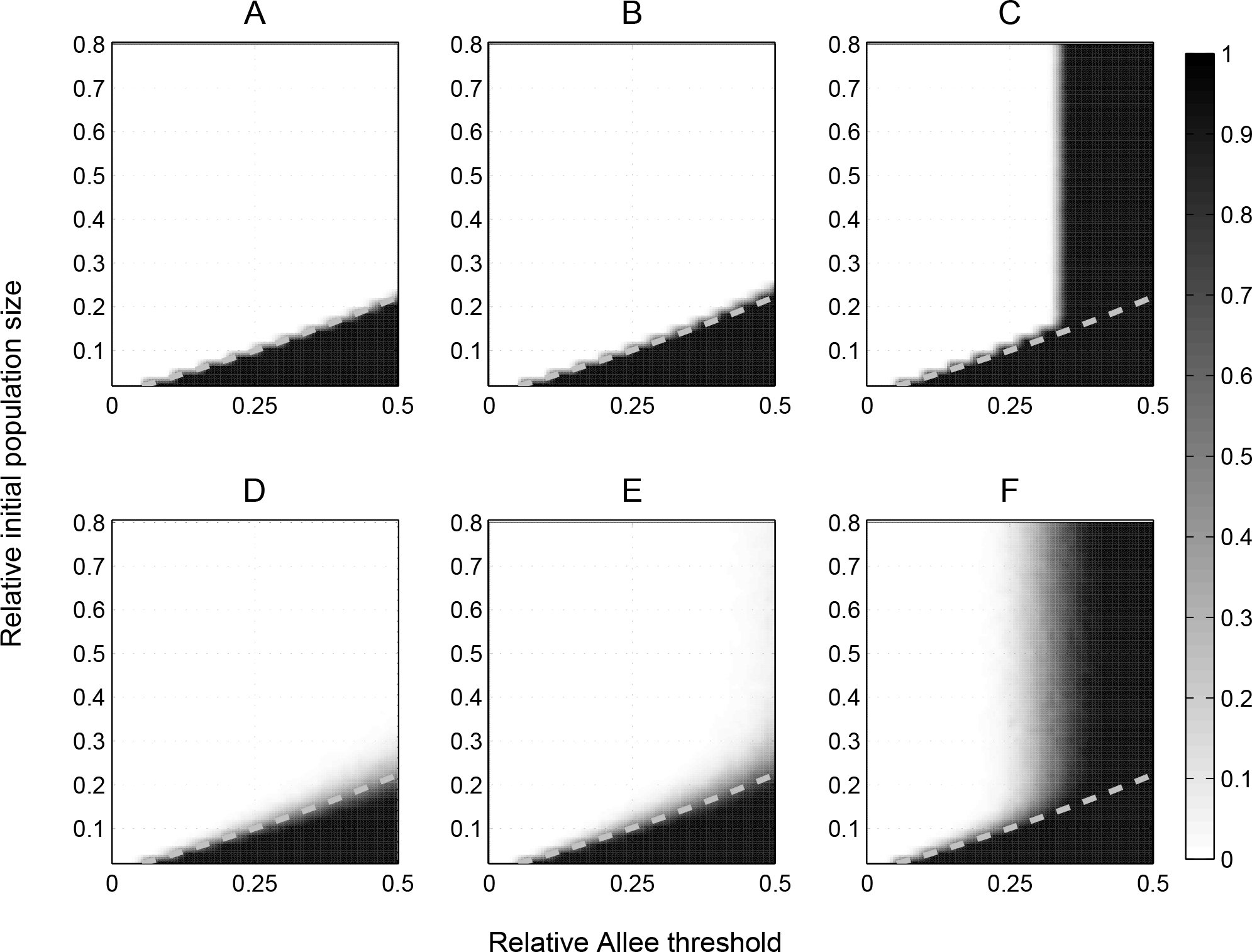
The gray scale in each plot represents the extinction probability for a given parameter combination based on 500 realizations of the model. In each plot, the initial population size and the strength of the Allee effect *θ*, both relative to *K*, are varied. We note that the carrying capacity of the population in the absence of Allee effects is 5500 with these parameters; therefore, the introduction sizes – which adjust the population size so that there is a 10% infection frequency ‣ do not exceed *∼* 612 insects. Top row: demographic stochasticity only. Bottom row: both demographic and environmental stochasticity. First column: no introduction. Second column: introduction such that infection frequency is at 10%. Third column: two subsequent introductions, both of which ensure the infection frequency is 10% for each cytotype at time of introduction (see Supplementary Information S2 for generation of second introduction). The dashed gray curve is the Allee threshold (i.e. initial populations below the gray curve go to extinction in the deterministic model). Here, *S*_0_ = 0.15.

In contrast to a single introduction, the release of two incompatible cytotypes is much more effective (Fig. 5C,F). Interestingly, when *θ* is relatively high, *Wolbachia* introductions succeed in driving population extinction independent of the initial population size. This result has the important implication that the success of *Wolbachia* introductions in driving extinction do not necessarily rely on having a pest population at the initial stage, or at the front, of the invasion. Moreover, in settings with high levels of environmental stochasticity, extinction is possible for much smaller Allee effects than predicted by the deterministic model (Fig. 5F).

As explored in the deterministic framework, combining *Wolbachia* introductions with other methods that increase the Allee threshold (e.g. mating disruption) will likely further increase the success of the overall management strategy. Here, we perform a more global exploration of management options that combine both *Wolbachia* introductions and pheromone releases. We again consider the population dynamics under three different management regimes: mating disruption only, mating disruption and the introduction of one cytotype, and mating disruption and the introduction of two cytotypes of *Wolbachia*. In this case, we assume that the relative strength of the Allee effect is fixed and relatively low so that *θ/K* is 0.1; under this condition, *Wolbachia* introductions alone do not drive the populations to extinction in the deterministic setting (Fig. 3B-C). We again find that environmental stochasticity increases the extinction probability for a smaller number of pheromone sources and, importantly, utilizing both mating disruption and CI is much more effective than using mating disruption alone (Fig. 6). Therefore, these two methods can serve as complementary tactics for pest management.

**Figure 6.**
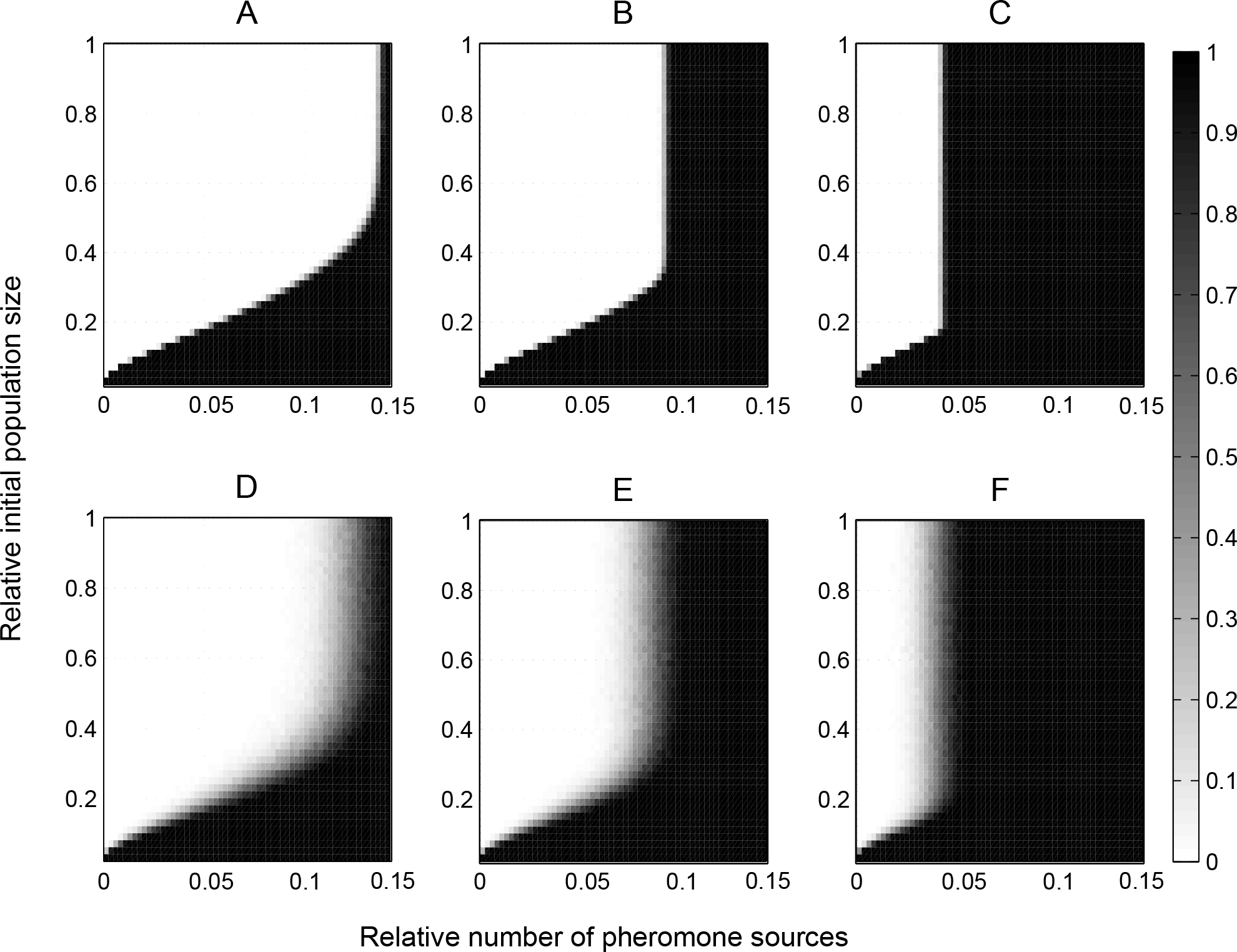
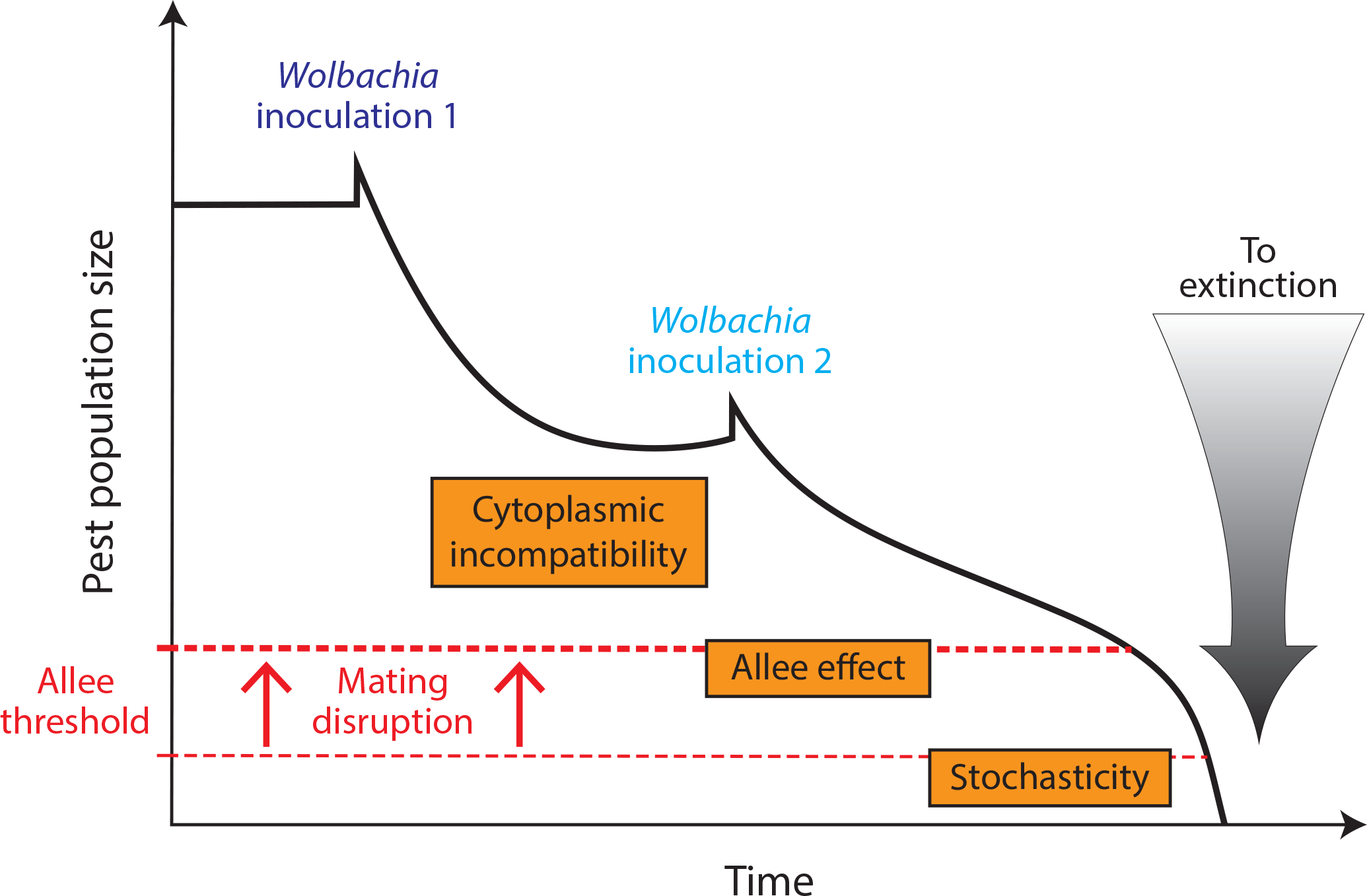
Fixing the relative strength of the Allee effect *θ/K* = 0.1, the gray scale in each plot represent the extinction probability for a given parameter combination based on 500 realizations of the model. In each plot, the initial population size and the number of pheromone sources (*P*) relative to *K* are varied. Top row: demographic stochasticity only. Bottom row: both demographic and environmental stochasticity. First column: no introduction. Second column: introduction such that infection frequency is at 10%. Third column: two subsequent introductions. Here, *S*_0_ = 0.15.

## Discussion

We investigated a population management strategy that considers *Wolbachia*-induced cytoplasmic incompatibility in the presence of Allee effects. In particular, we developed a stochastic population model, building upon the seminal approach of Dobson *et al.* (2002) and the continuously expanding body of literature investigating the use of Allee effects for the eradication of pest species (Liebhold & Bascompte, 2003; Tobin *et al.*, 2011; Liebhold *et al.*, 2016). Our model demonstrates that the introduction of a relatively small number of incompatible individuals into a pest population that has a strong pre-existing Allee effect can drive the pest population to extinction with no further intervention. These methods are successful more broadly when multiple strains of *Wolbachia* are introduced. We also demonstrate that extinction is possible for surprisingly large pest populations and that combinations of more than one strain of *Wolbachia* and mating disruption via sex pheromones work synergistically to increase the population’s extinction risk. Biological control has been studied for decades as an environmentally friendly alternative to pesticide use (e.g. Knipling, 1955; Baumhover *et al.*, 1955; Murdoch *et al.*, 1985; Bale *et al.*, 2008), and our study adds to this work by providing insight into ways that *Wolbachia* invasions can take advantage of intrinsic population processes – that is, Allee effects – to manipulate and control pest populations.

An important first step of our modeling work was to uncover the basic interactions between Allee effects and cytoplasmic incompatibility (CI). We show that these interactions are weak or non-existent: the *Wolbachia* invasion threshold does not depend on the strength of the Allee effect of its insect host, and the Allee threshold has only a marginal decrease in the presence of CI. Therefore, invasion of a particular *Wolbachia* strain into a population only depends on the critical infection frequency above which invasion succeeds in a deterministic setting (Barton & Turelli, 2011). This invasion threshold corresponds to a proportion of infected hosts above which infection spreads up to almost fixation, and is determined by parameters such as the reduction in egg hatch-rate caused by CI, the fitness costs of *Wolbachia* carriage, and the fraction of offspring that inherit the bacteria from an infected mother (Turelli, 1994). The invasion threshold found with our simulation model is consistent with that derived analytically (Turelli & Hoffmann, 1991), and unaffected by the intensity of a mate-finding Allee effect. In addition to adding validation to our model, this result holds interest because most theoretical approaches of *Wolbachia* invasion dynamics are purely genetic and consider changes in infection frequency without considering host population dynamics. One exception is the work of Hancock *et al.* (2011) which suggested that *Wolbachia* invasion thresholds predicted analytically hold for closed populations. Our results are consistent even when, as assumed here, host reproductive rate is affected by both positive and negative density dependence.

In the presence of strong Allee effects a population below the Allee threshold will be forced to extinction in a deterministic setting, making the Allee effect a central paradigm for conservation (Deredec & Courchamp, 2007; Stephens & Sutherland, 1999), invasions (Taylor & Hastings, 2005), biological control introduction (Fauvergue *et al.*, 2007, 2012), and as hypothesized in the present work, eradication (Tobin *et al.*, 2011). Whether an Allee effect is weak or strong (and the value of the Allee threshold in the latter case) depends on the strength of the underlying component Allee effect(s) relative to other density-dependent processes. Our simulations of various levels of cytoplasmic incompatibilities in a population with a pre-existing mate finding Allee effect suggest that the Allee threshold is much less sensitive to variations in the frequency of *Wolbachia*-infected individuals (0-90% infected individuals) than it is to variations in mating success (Fig. 2B). *Wolbachia*-induced cytoplasmic incompatibility does decrease population growth rate, as expected, but it has a minimal effect on the extinction threshold. Therefore, *Wolbachia*-induced CI may be considered a culling population management tactic where population size is temporarily decreased as a result of cytoplasmic incompatibilities.

Despite their initial apparent independence, cytoplasmic incompatibility and the Allee effect yield interesting properties when acting in concert. Our first analysis that considered the combined occurrence of Allee effects and CI in a deterministic context reveals that the transient decrease in population size is large enough to trigger extinction when the reproductive rate of the host species is relatively low. Extinction caused by the introduction of a single *Wolbachia* cytotype in populations with higher reproductive rates is only observed for very strong Allee effects (Fig. 3A). However, the strength of the Allee effect required for extinction lowers with the introduction of an additional incompatible *Wolbachia* strain. The resulting insect extinction probability, estimated by simulating the model in the presence of stochasticity, confirmed the interaction between the two processes. In the absence of *Wolbachia*, we determined the population’s extinction probability as it varies with its population size and the strength of the component Allee effect (Fig 5A and 5D). Introducing infected individuals results in the extinction of populations that would have persisted otherwise (i.e., a population that is above its Allee threshold can be brought to extinction). Introducing a second incompatible cytotype of *Wolbachia* increases CIs within the population and, consequently, increases the extinction domain by reducing the severity of Allee effect necessary to trigger extinction (Fig 5).

Nonetheless, our model predicts that although Allee effects and CI combine to drive populations to extinction – even surprisingly large populations – these extinctions may occur for unrealistically severe Allee effects. For instance, after the introduction of two incompatible *Wolbachia* strains into a population with a maximum reproductive rate of 3.75, extinction is expected when *θ/K* exceeds 0.25 (Fig. 5C-F); that is, extinction occurs if only 25% of all females successfully mate when the population is at 50% of the carrying capacity. Field estimations of mate-finding Allee effects in insects are rare, but it is probable that mating failures only occur at low densities. For instance, in the Gypsy moth *Lymantria dispar*, mating failures occurred below a density of *∼*4 (estimated via the rate of male captures on sex-pheromone traps) whereas the carrying capacity was estimated similarly around 800 (Tobin *et al.*, 2007, 2013), so that estimation of *θ/K* in this species could be one or two orders of magnitude lower than that yielding extinction in our model.

Our simulations demonstrate that eradication is much more likely if the introduction of *Wolbachia*-infected individuals is combined with mating disruption via the release of sex pheromone sources (Fig. 6). Interestingly, eradication is not restricted to small populations, but also applies to populations that have reached carrying capacity. Moreover, our results suggest that the two tactics act synergistically: the decrease in population size obtained when CI and mating disruption are combined is higher than the cumulative decrease obtained with each tactic applied separately. Our model therefore supports previous studies that highlight the potential benefit of simultaneously using multiple complementary management tactics (Blackwood *et al.*, 2012; Suckling *et al.*, 2012; Berec *et al.*, 2016). If different tactics benefit from one another, additional methods for controlling a pest population should also be considered. For example, other methods for population control such as parasitism or predation by native natural enemies may also be complementary. Additionally, while our focus was on cytoplasmic incompatibilities, there is evidence that *Wolbachia* and other bacteria are capable of other reproductive manipulations including male-killing (Dyer, 2004; Richardson *et al.*, 2016). Similar conclusions were also made in the recent modeling study of Berec *et al.* (2016), who suggest that sterile insect technique is improved when combined with male-killing bacteria. This suggests the existence of additional avenues for utilizing *Wolbachia* in the context of pest management.

Theoretical models of population management are currently flourishing much faster than empirical evidence can be obtained; therefore, it is important to discuss the relevance of our predictions and the feasibility of our proposed methods. First, we have shown that extinction in insect species with high reproductive rates may not be feasible because extinction would require an unrealistic amount of sex pheromone lures to successfully complement the *Wolbachia*-induced transient decrease in population size. However, in the majority of our simulations we assigned the maximum reproductive rate a value of 3.75 or, equivalently, *per capita* fecundity as *m* = 25 and density independent survivorship of larvae as *S*_0_ = 0.15. This value is just below the median of the reproductive rates estimated in Hassel *et al.* (1976) across 24 different insect species. Therefore, we conjecture that these methods would still apply to a variety of pest species. Second, the reproductive rate is just one of the parameters in our model. Although the main text is supplemented with sensitivity analyses, our work is not intended to provide robust quantitative guidelines for a practical application of our proposed management strategies. Instead, our analysis provides general properties of the interactions between Allee effects and cytoplasmic incompatibilities on the population dynamics of an insect pest species. Should our proposed management strategy be applied to a specific insect pest that exhibits Allee effects, thorough investigations would be needed to better quantify parameters that dictate the strength of the processes that underpin the model, including *Wolbachia* diversity and incompatibility as well as component and demographic Allee effects.

To date, the potential use of *Wolbachia*-induced CI for pest management is supported by a few but important studies on mosquitoes (Hoffmann *et al.*, 2011; Laven, 1967) and fruit flies (Zabalou *et al.*, 2004). Furthermore, evidence for the existence of both a component and a strong demographic Allee effect exists in the Gypsy moth, which could explain the relative success of mating disruption in this species (Tobin *et al.*, 2013, 2007). Although such empirical advances are indeed promising, they deserve a much stronger body of data and robust cause and effect demonstrations (Fauvergue, 2013). From this perspective, our model was not built as a predictive tool for a specific species. It was rather developed as a heuristic theory yielding qualitative predictions which will hopefully encourage future experimental approaches on the consequences of cytoplasmic incompatibility and Allee effects on population extinction.

There is a long and prolific body of research in population dynamics that focuses on understanding the mechanisms stabilizing species near their carrying capacities (e.g. Hassell & May, 1973; Robert M. May, 1978; Bernstein, 2000). More recently, global climate change and the biodiversity crisis, including population declines, extinctions, or biological invasions, points towards the increasing relevance of nonequilibrium ecology (Rohde, 2006) and the biology of small populations (Fauvergue *et al.*, 2012). Transient dynamics are increasingly emphasized (Hastings, 2004) and sometimes considered in the specific context of population management (Ezard *et al.*, 2010; Kidd & Amarasekare, 2012). As first highlighted by Dobson *et al.* (2002), cytotype replacement which occurs in the course a successful *Wolbachia* invasion yields a transient coexistence of incompatible infections within a host population, and as a consequence, a transient decrease in reproductive rate and population size. Here, the transients only last a few generations and this perturbation of the population’s microbiome is the first step in a destabilizing cascade. We show here that the population can then be pushed toward a second step of destabilization, triggered by a mate-finding Allee effect that can be reinforced by the application of mating disruption, which potentially drives the population to extinction.

## Acknowledgements

We thank Fabrice Vavre for his enthusiasm in preliminary discussions, and Sylvain Charlat for comments in earlier stages of this work. We also thank the Associate Editor and two anonymous reviewers for their helpful and constructive feedback.

